# Cerebellar and Prefrontal-Cortical Engagement During Higher-Order Rule Learning in Older Adulthood

**DOI:** 10.1101/2020.01.22.914739

**Authors:** T. Bryan Jackson, Ted Maldonado, Sydney M. Eakin, Joseph M. Orr, Jessica A. Bernard

## Abstract

To date most aging research has focused on cortical systems and networks, ignoring the cerebellum which has been implicated in both cognitive and motor function. Critically, older adults (OA) show marked differences in cerebellar volume and functional networks, suggesting it may play a key role in the behavioral differences observed in advanced age. OA may be less able to recruit cerebellar resources due to network and structural differences. Here, 26 young adults (YA) and 25 OA performed a second-order learning task, known to activate the cerebellum in the fMRI environment. Behavioral results indicated that YA performed significantly better and learned more quickly compared to OA. Functional imaging detailed robust parietal and cerebellar activity during learning (compared to control) blocks within each group. OA showed increased activity (relative to YA) in the left inferior parietal lobe in response to instruction cues during learning (compared to control); whereas, YA showed increased activity (relative to OA) in the left anterior cingulate to feedback cues during learning, potentially explaining age-related performance differences. Visual interpretation of effect size maps showed more bilateral posterior cerebellar activation in OA compared to YA during learning blocks, but early learning showed widespread cerebellar activation in YA compared to OA. There were qualitatively large age-related differences in cerebellar recruitment in terms of effect sizes, yet no statistical difference. These findings serve to further elucidate age-related differences and similarities in cerebellar and cortical brain function and implicate the cerebellum and its networks as regions of interest in aging research.

## Introduction

Even in healthy aging, older adults (OA) experience declines in both cognitive and motor performance. Many authors have reported age differences in fluid cognitive processing (Anguera, Reuter-Lorenz, Willingham, & Seidler, 2011; Bo, Jennett, & Seidler, 2012; Deary et al., 2009; Harada, Love, & Triebel, 2013; Manard, Carabin, Jaspar, & Collette, 2014) and motor behaviors (Ketcham, Seidler, Van Gemmert, & Stelmach, 2002; Seidler, Alberts, & Stelmach, 2002; Verghese et al., 2002), each of which have been linked to differences in brain structure and functional activation patterns (reviewed in Reuter-Lorenz & Lustig, 2005; Seidler et al., 2010). Currently, 9% of the world’s population are aged 65 and older, and by 2050, OA will comprise 17% of the population (Roberts, Ogunwole, Blakeslee, & Rabe, 2018). Given the dramatic increase in this demographic, differences in brain structure and function as they relate to performance is of the utmost importance, as this provides a key first step to improving quality of life and health for aging individuals.

Age-related neuroanatomical differences have been shown in both grey matter and white matter. Structurally, many brain regions show smaller volume reduction across tissue types (Abe et al., 2008; Giorgio et al., 2010; Seidler et al., 2010), increased white-matter hyperintensities (de Leeuw, 2001), and a reduction in white-matter integrity with age (Maniega et al., 2015). Additionally, differences in cortical functional connectivity and resting state networks are also seen in OA. In healthy young adults (YA), task-positive networks and the default mode network (DMN) are anticorrelated (Fox et al., 2005; Spreng, Stevens, Viviano, & Schacter, 2016); however, prior work suggests OA show reduced task-related suppression of the DMN and changes in activation of task-positive networks (Grady et al., 2010; Ng, Lo, Lim, Chee, & Zhou, 2016; Spreng et al., 2016).

Motor and cognitive task-related cortical functional activation also exhibits age-related differences. For example, picture recognition (Morcom, Li, & Rugg, 2007), word recognition (Duverne, Motamedinia, & Rugg, 2009), and working memory paradigms (Emery, Heaven, Paxton, & Braver, 2008) indicate that OA display increased activation in task positive regions, compared to performance-matched YA. OA also exhibit increased activation and lower absolute grip force compared to YA when relative grip force is matched between groups (Noble, Eng, Kokotilo, & Boyd, 2011). Models such as the Compensation-Related Utilization of Neural Circuits Hypothesis (CRUNCH; Park & Reuter-Lorenz, 2008; Reuter-Lorenz & Lustig, 2005) and Hemispheric Asymmetry Reduction in Older Adults (HAROLD; Cabeza, 2002; Cabeza, Anderson, Locantore, & McIntosh, 2002) conceptualized age-related brain activation differences as compensatory scaffolding for decreased processing efficiency. Taken together, research suggests that age-related performance differences may be partially related to changes in task-related brain activation and connectivity between brain regions, in conjunction with changes in brain structure seen in advanced age.

CRUNCH (Park & Reuter-Lorenz, 2008; Reuter-Lorenz & Lustig, 2005) and HAROLD (Cabeza, 2002; Cabeza et al., 2002), along with other accounts of age differences in functional activation, have provided important insights into brain-behavior relationships in advanced age. But, the primary focus of these frameworks is cortical, and primarily prefrontal cortical at that. A robust and growing literature implicates long-range networks of the cerebellum in cognitive processing. Indeed, cerebellar activity is frequently noted during cognitive tasks in YA (Chen & Desmond, 2005; Guell, Gabrieli, & Schmahmann, 2018; for review see Schmahmann, 2019), and both non-human primate (Dum & Strick, 2003; Kelly & Strick, 2003; Strick, Dum, & Fiez, 2009) and human research (Bernard, Orr, & Mittal, 2016; Bernard et al., 2012; Krienen & Buckner, 2009; O’Reilly, Beckmann, Tomassini, Ramnani, & Johansen-Berg, 2010; Salmi et al., 2010) has shown evidence of distinct motor and cognitive loops, separately connecting the cerebellum to the primary motor cortex and prefrontal cortex via the thalamus (cerebello-thalamo-cortical circuits; CTCC). Further, a functional topography within the cerebellum has been described that closely parallels these closed-loop circuits (King, Hernandez-Castillo, Poldrack, Ivry, & Diedrichsen, 2019; Stoodley & Schmahmann, 2009; Stoodley, Valera, & Schmahmann, 2012). Given the cortical differences in older adults and evidence of cerebellar recruitment during cognition in YA, it is of interest to investigate whether and how the cerebellum contributes to cognitive processing in advanced age. Indeed, this subcortical structure may provide key scaffolding for cognition, as has been suggested in motor tasks (Filip, Gallea, Lehéricy, Lungu, & Bareš, 2019). Critically, there is evidence that the cerebellum and its resting state networks are negatively impacted with age in behaviorally relevant ways (smaller volume, decreased connectivity; Abe et al., 2008; Bernard et al., 2013; Bernard & Seidler, 2014; Hausman, Jackson, Goen, & Bernard, 2019), further underscoring the importance of investigating cerebellar function in advanced age.

To this point, targeted investigations of age differences in cerebellar functional activation are relatively lacking. A recent report by Filip and colleagues (2019) detailed bilateral posterior cerebellar hyperactivation that increased with age during a predictive motor timing task, suggesting compensatory neural scaffolding comparable to patterns found in the prefrontal cortices (Park & Reuter-Lorenz, 2008). Conversely, under-activation of lobule HVI, a cerebellar region associated with motor learning, was found during a symbolic motor sequence learning task (Bo, Peltier, Noll, & Seidler, 2011), consistent with predictions by Bernard & Seidler (2014). It is therefore an open question as to whether task-related cerebellar activity mirrors the overactivation patterns found in the prefrontal regions or is hindered by decreased communication between the two regions, as proposed by Bernard and Seidler (2014).

Here, we used whole-brain functional imaging to investigate age differences in task performance and functional activation patterns in OA and YA to provide broader insights into brain function in advanced age. We adapted an in-scanner second-order rule learning task (Balsters, Whelan, Robertson, & Ramnani, 2013), known to elicit activation of the posterior lobules of the cerebellum in healthy YA and previously used by our lab in a clinical investigation (Bernard, Orr, Dean, & Mittal, 2018). Consistent with visuomotor and higher-order learning performance in OA (Anguera et al., 2011; Howard & Howard, 1997; Seidler, 2006), we hypothesized that performance differences would be found in advanced age, evidenced by lower accuracy and a slower rate of learning of second-order rules. In addition, we planned a group contrast to compare functional activation patterns as they relate to higher-order rule learning during the preparation and response period of the task. Due to the conflicting nature of the literature, we predicted that OA would show either compensatory bilateral overactivation of posterior cerebellar lobules, consistent with what has been shown in the prefrontal cortex and in the cerebellum during predictive motor timing (Filip et al., 2019; Park & Reuter-Lorenz, 2008), or under-activation, as was found for motoric regions of the cerebellum during a serial reaction task (Bo et al., 2011) and consistent with a recent review on the topic (Bernard & Seidler, 2014).

## Methods

### Participants

Here, 25 OA (mean age = 72.24 ± 6.29; age range = 60-84; 16 females) and 26 YA (mean age = 22.54 ± 2.87; age range = 18-30; 14 females) were recruited from the Bryan-College Station, Texas community and compensated for their time. One participant from each group was left-handed (3.9%), notably below the proportion of left-handed individuals in the population (approximately 10%; Oldfield, 1971). All participants were cognitively healthy, as confirmed using the Montreal Cognitive Assessment (MoCA; Nasreddine et al., 2005). Exclusionary criteria for both groups consisted of a history or stroke or neurological disease and contraindications for the neuroimaging environment. One OA participant performed poorly throughout the learning blocks (<5% correct) and was removed from all analyses, resulting in a final sample of 24 OA and 26 YA. Prior to beginning study procedures, all participants provided their informed consent. All study procedures were approved by Texas A&M University’s Institutional Review Board and in compliance with the Helsinki Declaration of 1975 (as revised in 1983).

### Second-order rule learning task

To assess non-motor rule learning and CTCC function, we used a paradigm developed by Balsters and colleagues (2013), that was recently adapted and implemented by our group (Bernard et al., 2018). This second-order rule learning task dissociates the learning of the rule itself from the motoric responses. Here participants completed 4 blocks of the learning task, and 2 control blocks. Each block consisted of 25 trials. In our recent work, we found that performance plateaus after 4 blocks in healthy YA (Bernard et al., 2018), and as such, we limited the task to 4 blocks here. Further, this prevents fatigue for the OA group.

During each trial, participants viewed one of four abstract shapes (“instruction cue”), presented in green on a gray background, for 500 ms. Then, after a variable delay (onset was jittered), the word “Go!” was presented in red on a gray background for 250 ms. This was immediately followed by a cue onscreen for 1000 ms indicating an answer should be provided (“reaction cue,”). The reaction cue consisted of four differently colored hourglasses (pink, blue, yellow, and black), with their position randomized between trials. Participants were to use trial and error to match each instruction cue to a preselected color hourglass. Feedback cues were given immediately in the form of a green circle for correct responses, a red circle for incorrect responses, and “Missed!”, presented in red, when a response was not received. The feedback cue was onscreen for 250 ms and all feedback was presented on a gray background. To accommodate the scanner environment, each trial was divided into two 4-second sections (2 TRs each): the instruction cue period (instruction cue and surrounding ISIs) and the reaction cue period (“Go!”, reaction cue, feedback, and surrounding ISIs), each jittered to alleviate metronomic effects.

In addition to the second-order rule learning task, participants also completed two control blocks in which the instruction cue was always the same shape (unique from those during the learning blocks) and the reaction cue consisted of only one hourglass of random color in one of four random locations. Participants were asked to respond to the location of the hourglass. This design allowed us to look at activation during the instruction cue of the learning blocks, presumably related to second-order rule learning, contrasted against a simplified, strictly motor version of the task without any learning elements. The instruction cue was of interest because the participants would be processing the second-order rule, deciding on the shape/color pair, as well as anticipating the reaction cue. The reaction cue was of interest because participants would be processing the second-order rule, deciding which spatial location contains the correctly colored hourglass, and responding. Finally, the feedback cue was of interest because it would trigger a rule update and allow us to investigate the processing in response to error information.

Finally, while in the scanner, participants also completed two learning blocks under dual-task conditions, before and after the learning task, to asses learning and automaticity. The dual-task blocks were formatted exactly like the learning blocks discussed above, with the additional requirement of having to verbally count backwards by 7 from 500. Though we did not record counting accuracy, participants were prompted to continue counting when no responses were made for >3 s. Imaging data was not recorded so that the verbal counting could be heard over the scanner noise and to avoid motion confounds due to speech. We quantified performance on these dual-task blocks and calculated the dual-task cost. The dual-task cost was calculated by subtracting accuracy of learning block 4 from post-test dual-task accuracy and represents the degree to which the rule and associations have been automatized. Positive values mean performance was better under single-task conditions, with higher values indicating a greater cost under dual-task conditions. The dual-task cost was compared between groups using an independent-samples t-test.

### Behavioral analysis

Demographic and behavioral data were analyzed using IBM SPSS 25 (IBM Corporation Armonk, NY, 2017). Independent-sample t-tests were used for group differences in demographics.

Accuracy was calculated separately for each block (pre, control 1 and 2, learning 1, 2, 3, and 4, post) using the percent of correct responses.

First, in order to verify that learning occurred, a 2 (YA, OA) x 2 (pre-test, post-test) repeated-measures mixed model ANOVA was used to compare pre- and post-test accuracy under dual-task conditions. Slopes were then calculated for each consecutive block of increasing length (e.g. block 1 through 2, blocks 1 through 3, and blocks 1 through 4) during the learning task to quantify the rate of learning. The rate of learning was analyzed by calculating the learning slope from block 1 to all other blocks for each group separately. Learning slopes were then compared using a 2 (OA, YA) x 3 (slope1-2, slope 1-3, slope 1-4) repeated-measures mixed model ANOVA and followed up with paired t-tests comparing OA and YA. The steepest learning slope for each group, for use in the ROI correlations described below, was determined by comparing subsequent slopes in a paired t-test within each age group. If statistically significant differences were seen between multiple slope calculations, the slope with the largest magnitude was used. For the sake of completeness, we also compared the accuracy of each learning block using a 2 (YA, OA) x 4 (blocks 1, 2, 3, 4) repeated measures mixed model ANOVA and followed up with paired *t*-tests comparing blocks across groups. The control blocks were analyzed using a 2 (YA, OA) x 2 (control block 1, control block 2) repeated measures mixed ANOVA and followed up using independent-samples *t*-tests comparing OA and YA.

### fMRI data acquisition

fMRI data were collected using a 3-tesla Siemens Verio scanner with a 32-channel head coil at the Texas A&M Translational Imaging Center. Blood-oxygen-level dependent (BOLD) scans were acquired in alternative phase encoding directions with an echo-planar functional protocol and a multiband factor of 4 (number of volumes =102, repetition time [TR]=2000 ms, echo time [TE]= 27 ms; flip angle [FA]=52°, 3.0×3.0×3.0 mm3 voxels; 56 slices, field of view (FOV)= 300 x 300 mm; time= 3:36 min). In addition, a high-resolution T1-weighted 3D magnetization prepared rapid gradient multi-echo sequences (MPRAGE; sagittal plane; TR=2400 ms; TE=2.07 ms; 0.8 mm3 isomorphic voxels; 160 slices; FOV=256 x 256 mm; FA=8°; time=7:02 minutes) to facilitate normalization of the fMRI data.

### fMRI data pre-processing and analysis

Anatomical and functional images were collected in DICOM format then converted to NIFTI files and organized into a Brain Imaging Data Structure (BIDS) compliant directory using the latest version of bidskit (version 1.1.2; Mike Tyszka 2016). Each functional image was collected in pairs of images with opposite phase encoding. In order to create pairs of opposite encoded single volumes for distortion correction, a single volume from two of each participant’s 4D functional images was extracted then combined. FSL’s topup utility was then used to generate fieldmap images for use in unwarping the images.

FMRI data was preprocessed using FEAT (FMRI Expert Analysis Tool) Version 6.00, part of FSL (FMRIB’s Software Library, www.fmrib.ox.ac.uk/fsl; Jenkinson, Beckmann, Behrens, Woolrich, & Smith, 2012). The following preprocessing methods were applied: motion correction using MCFLIRT (Jenkinson, Bannister, Brady, & Smith, 2002); slice-timing correction using Fourier-space time-series phase-shifting; non-brain removal using BET (Smith, 2002); spatial smoothing using a Gaussian kernel of FWHM 6.0mm; grand-mean intensity normalization of the entire 4D dataset by a single multiplicative factor; highpass temporal filtering (Gaussian-weighted least-squares straight line fitting, with sigma=45.0s). ICA-based exploratory data analysis was carried out using MELODIC (Beckmann & Smith, 2004), in order to apply automated ICA-based denoising using FIX (described in more detail below). Registration to high resolution structural and/or standard space images was carried out using FLIRT (Jenkinson et al., 2002; Jenkinson & Smith, 2001). Registration from high resolution structural to standard space was then further refined using FNIRT nonlinear registration (Andersson, Jenkinson, & Smith, 2007).

40 blocks, each from a different subject, were chosen at random (10 YA and 10 OA control blocks and 10 YA and 10 OA learning blocks) and their ICA components were visually inspected to determine which components represented noise or artifacts (e.g., pulse, breathing, motion, multiband artifacts, etc.). FIX (v 1.0.68; Griffanti et al., 2014; Salimi-Khorshidi et al., 2014), an automatic component classifier, was used to create 4 separate training files (1 each for OA control blocks, YA control blocks, OA learning blocks, and YA learning blocks), which were subsequently used to remove noise and artifacts from the entire sample’s functional data prior to statistical analysis. Time-series statistical analysis was carried out using FILM with local autocorrelation correction (Woolrich, Ripley, Brady, & Smith, 2001). All within-subject variables were modeled as fixed effects and group contrasts were modeled using FLAME 1+ 2 (FMRIB’s Local Analysis of Mixed Effects). FIX was performed with an Apple iMac with a 3.1GHz Intel Core i5 processor; all other preprocessing and whole brain analyses were accomplishing using a local distributed computer cluster (Brazos Computational Resource, Texas A&M University).

Activity was time-locked to the onset and duration of each instruction cue, reaction cue, and feedback cue. Only correct responses were included in our analyses of the instruction cue and reaction cue. Our interest in learning and updating led us to include all activity during feedback cues for two reasons. First, negative feedback is important in these processes. Second, because YA learned the task very quickly, there were not enough incorrect responses to have the power to investigate correct and incorrect feedback separately. We modeled two within-subjects contrasts. We collapsed across the four learning blocks and contrasted the mean against the mean of the control blocks to compare task-related activation in the two age groups. Since we were interested in learning and YA had learned the correct responses by block 2, blocks 1 and 4 were contrasted (separately from the control blocks) to investigate areas activated early during the learning process with those activated later in the process. Individual group means and age contrasts were modeled and thresholded nonparametrically using clusters determined by *z* = 3.1, and a cluster significance threshold of *p_fwe_* < .05 using non-parametric methods with 5000 permutations (Winkler, Ridgway, Webster, Smith, & Nichols, 2014). A *post-hoc* investigation exploring overall patterns of activation by collapsing across groups was performed using the same statistical parameters.

To further detail and capture subthreshold age-related differences in cerebellar functional activation, exploratory effect size calculations were performed using OA > YA and YA > OA contrast images for all three cues. Group contrast *t*-stat maps used in the primary analyses were converted to Cohen’s *d* effect size maps using fslmaths and thresholded at *d* = .5 using fsl’s cluster command (medium effect size; Cohen, 1970). Images are presented for visual interpretation.

## Results

### Behavioral performance

Table 1 lists the means and standard deviations for accuracy, separately for each block and condition, slopes from block 1 for each subsequent block, and dual task cost.

**Table 1.**
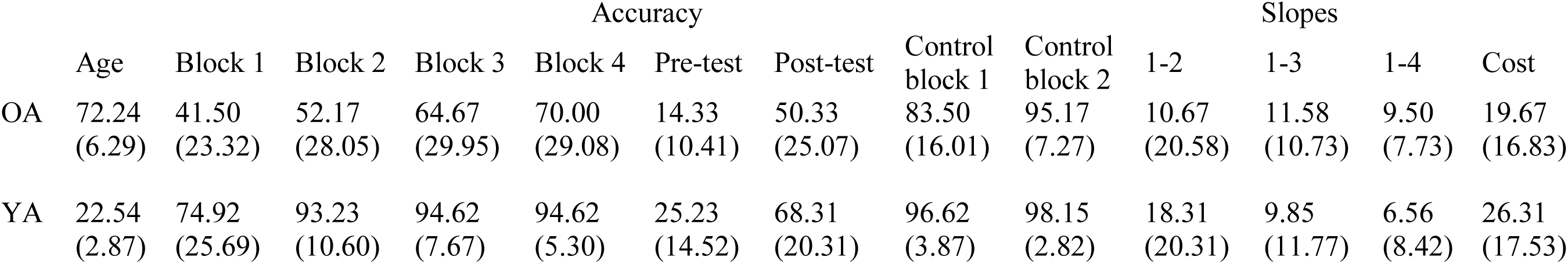
Means (and standard deviations) for age and dependent variables, separately by group.

#### Dual Task

Learning was verified by analyzing dual-task performance. A main effect of task (post-test > pre-test, *F*(1, 48) = 148.49, *p* < .001, 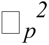 = .76) and of group (YA > OA, *F*(1, 48) = 148.06, *p* < .001, 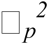 = .90) was found (Figure 1A). There was no significant interaction (*F*(1, 48) = 1.19, *p* = .28, 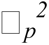 = .02). We also investigated group differences in dual-task cost after learning and found no difference between groups (*t*(48) = 1.36, *p* = .18, *d* = .39) (Figure 1B).

**Fig. 1.**
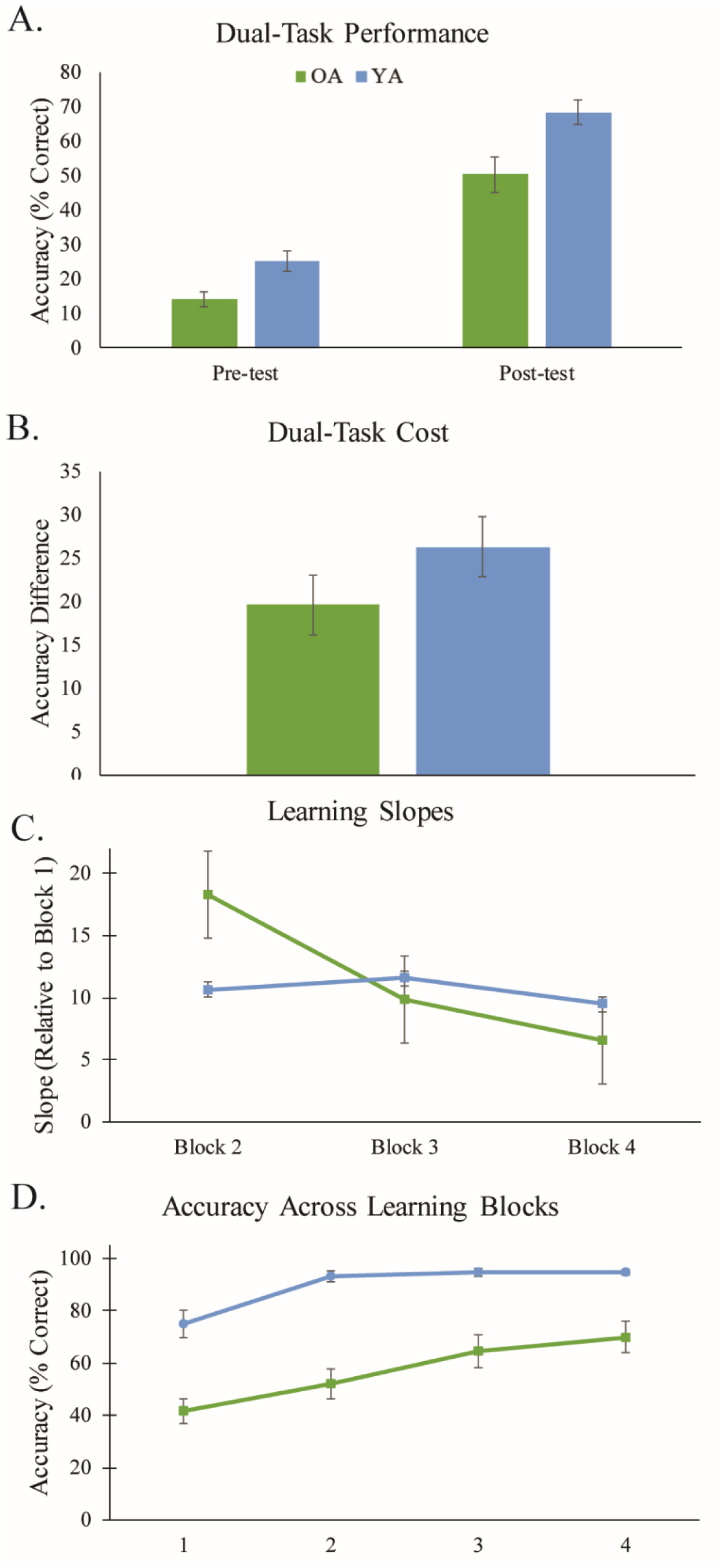
Depictions of behavioral results comparing OA (green) to YA (blue). A) Bar graph depicting pre- and post-test accuracy averages. B) Bar graph depicts the dual-task cost between learning block 4 and the post-test average. C) Line chart depicting average incremental slope for each age group. D) Line chart depicting average accuracy for each learning block.

#### Learning Slopes

There was a significant group X learning slope interaction (*F*(2, 96) = 5.67, *p* = .01, 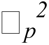 = .11), and follow up paired *t*-tests indicated significant differences between all slopes in YA, with each progressive slope smaller than the last (slope 1-2 versus slope 1-3, *t*(25) = 4.47, *p* < .001; slope 1-2 versus slope 1-4, *t*(25) = 4.73, *p* < .001; slope 1-3 versus slope 1-4, *t*(25) = 4.07, *p* < .001). This indicates that YA showed the fastest rate of learning during the first two blocks, reaching ceiling performance thereafter; whereas, OA showed no difference between any slopes (*ps* > .12), indicating a more consistent and slower rate of learning. Main effects of slope (*F*(2, 96) = 7.11, *p* = .001, 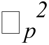 = .129) and group (YA>OA: *F*(1, 48) = 39.68, *p* <.001, 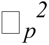 = .45) were also found (Figure 1C).

#### Accuracy

For control blocks, there was a main effect of group (*F*(1, 48) = 8382.42, *p* < .001, 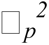 = .99) as well as a block x group interaction (*F*(1, 48) = 11.36, *p* = .001, 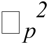 = .019) indicating that OA performed lower on control block 1 (*t*(48) = 4.05, *p* < .001, *d*= 1.13), but did not differ for control block 2 (*t*(48) = 1.94, *p* = .07, *d* = .54).

During the learning task, results indicated a block x group interaction (*F*(3, 144) = 3.41, *p* = .02, 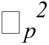 = .07), indicating OA performed worse and learned more slowly than YA (Figure 1D). Main effects of block (*F*(3, 144) = 33.40, *p* < .001, 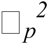 = .41) and group were also found (YA>OA: *F*(1, 48) = 49.99, *p* < .001, 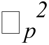 = .51).To limit multiple comparisons, further accuracy analyses were forgone, and we focused entirely on the slope (learning rate) analysis presented here.

### Brain activation patterns

Tables 2-4 detail the anatomical region, Broadmann’s Area (BA), size (in voxels), *t*-statistic, Montreal Neurological Institute (MNI) space coordinates, and effect size (Cohen’s *d*) of each activated cluster’s peak. Local maxima are listed in italics. Anatomical regions from the Automated Anatomical Labeling atlas and Brodmann Areas were determined using label4MRI (Chuang and Yun-Shiuan, 2019; R package version 1.2).

**Table 2.**
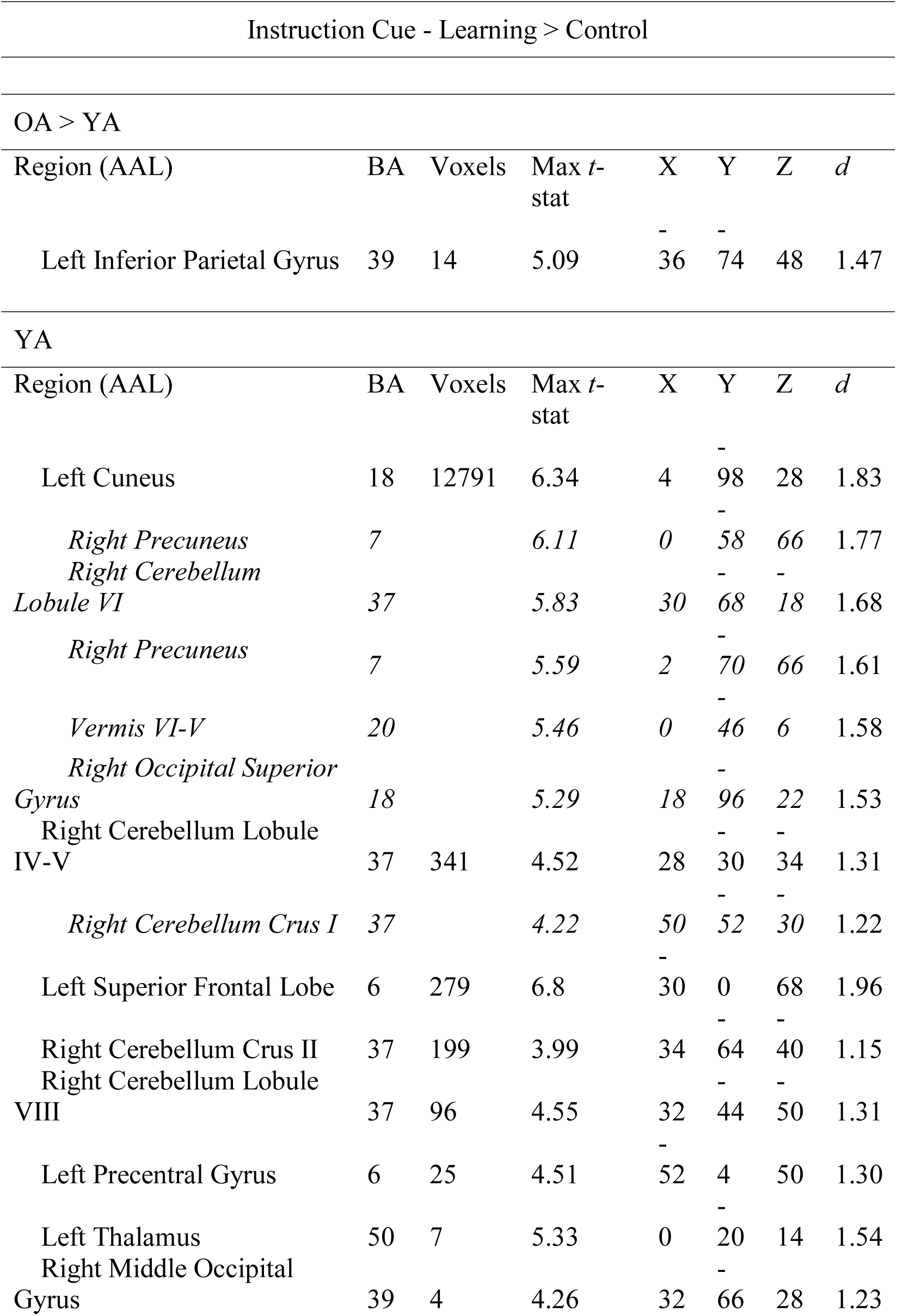
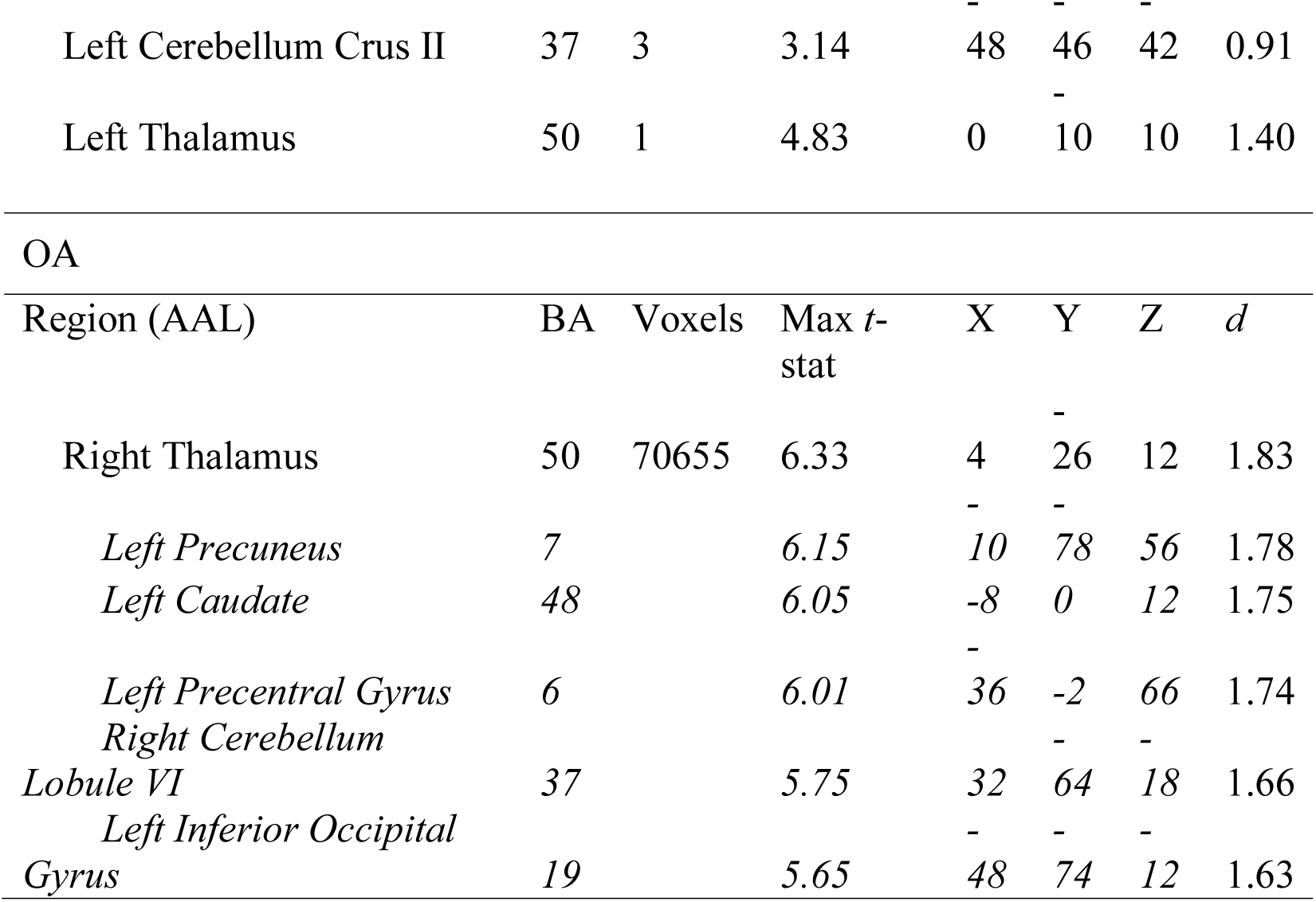
Areas of peak activation in response to instruction cues on correct trials during all learning blocks compared to control blocks. Groups differences and within group data are presented. Regions in italics are local maxima within a larger cluster.

#### Learning > Control

Table 2 details activation patterns to instruction cues in the group contrast and within group images, for learning compared to control blocks. During instruction cue presentation, OA showed greater activation than YA in the left inferior parietal lobe (Figure 2A). YA did not have any activations larger than OA. Within OA alone, a large cluster showed activation throughout subcortical regions and the cerebello-thalamo-frontal network; much of the posterior cerebellum was active, with a local peak near lobule VI (Table 2; Supplemental Figure 1). YA showed widespread activation, including right cerebellum Crus I and II, bilateral thalamic activation, right precuneus, left cuneus, and left superior frontal and precentral gyri (Table 2; Supplemental Figure 2). Effect size calculations indicated that OA showed more activation compared to YA in the posterior cerebellum (Figure 3A), with medium-to-large effect sizes, while YA had almost no cerebellar activation greater than OA (Figure 3B).

**Fig. 2.**
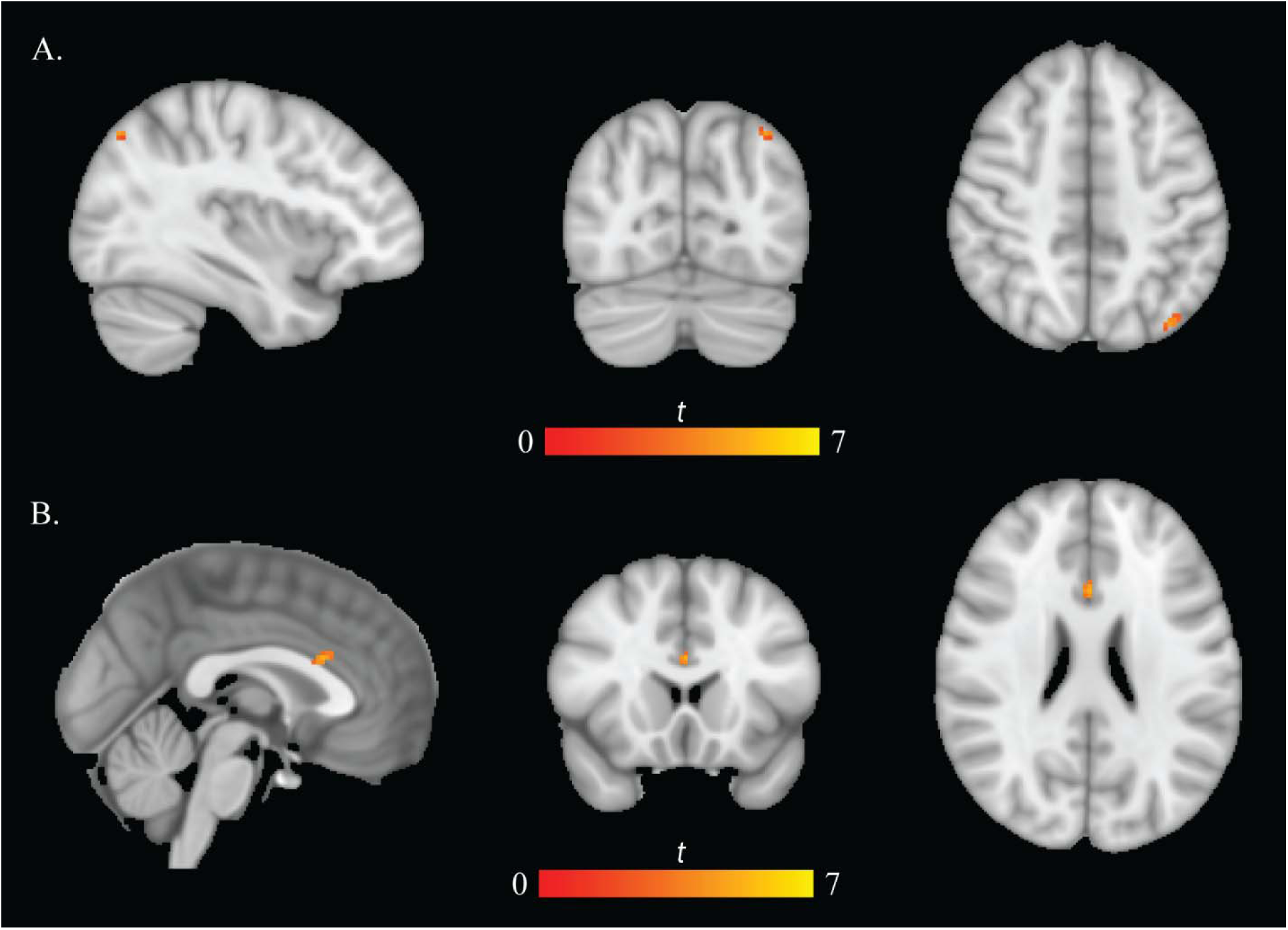
A) Brain activation patterns to instruction cues during learning. Left inferior parietal lobe shows more activation in OA than YA. B) Brain activation patterns to feedback cues during learning. Left anterior cingulate shows more activation in YA than OA.

**Fig. 3.**
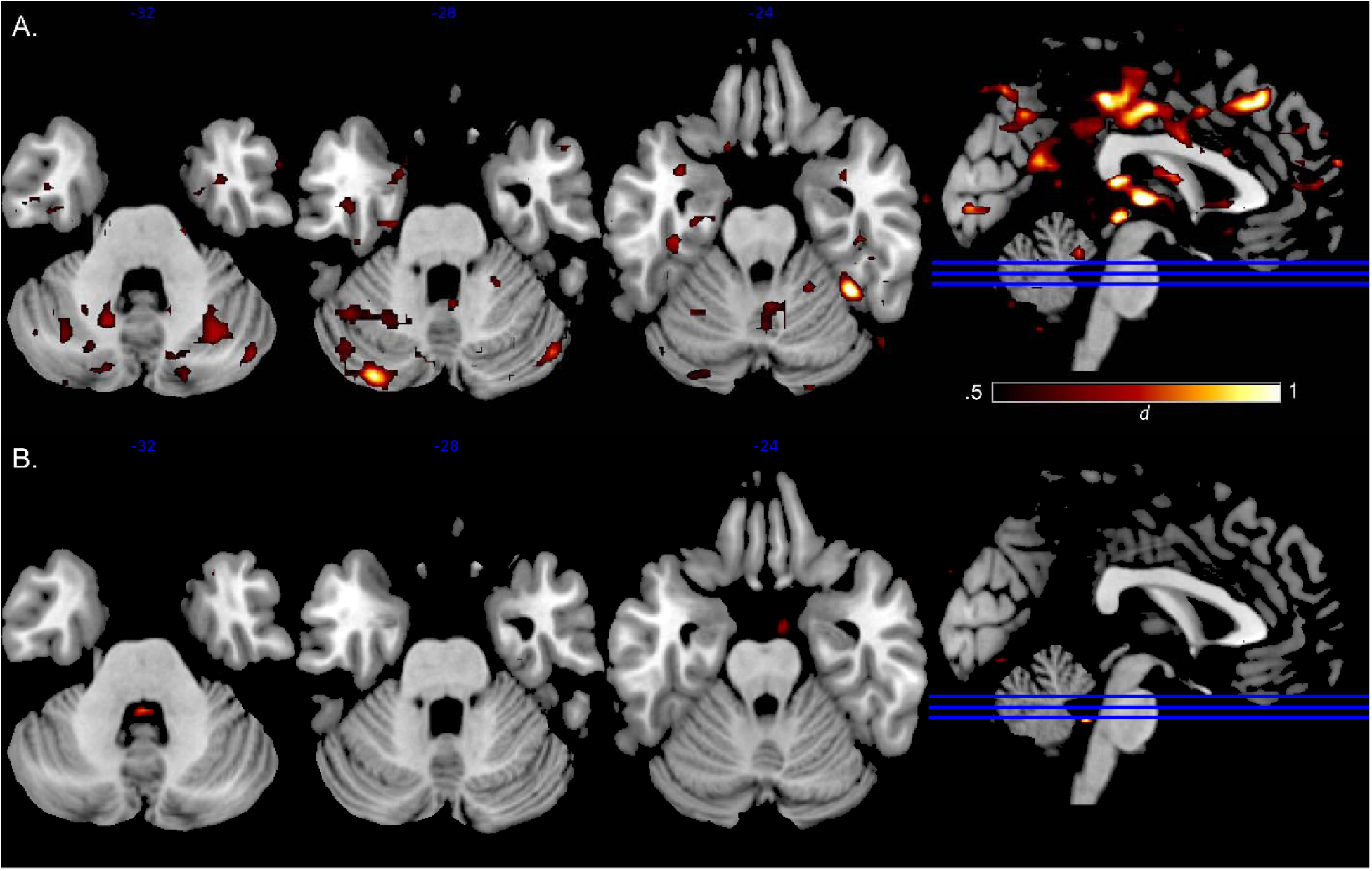
Effect size maps to instruction cues during learning for (A) OA > YA and (B) YA > OA.

No significant clusters were found in response to reaction cues in the learning > control contrast.

Table 3 details activation patterns to feedback cues in the group contrast and within group images, for learning compared to control blocks. Feedback cues elicited greater activation in the left anterior cingulate in YA compared to OA (Table 3; Figure 2B). OA did not show any greater activation compared to YA. Within OA, multiple bilateral regions in the frontal and temporal lobes were activated, as well as the bilateral precentral gyrus, left cerebellum Crus II, and left middle cingulate (Table 3; Figure 4). YA showed activation in frontal lobes (mostly contained within the right hemisphere), bilateral anterior cingulate, right precentral gyrus, bilateral precuneus, bilateral cerebellar Crus I, and the right insula (Table 3; Figure 5). Effect size calculations indicated that OA showed more activation compared to YA in the posterior cerebellum, with medium-to-large effect sizes; whereas, YA (compared to OA) showed very little posterior cerebellar activation (Figure 6).

**Fig. 4.**
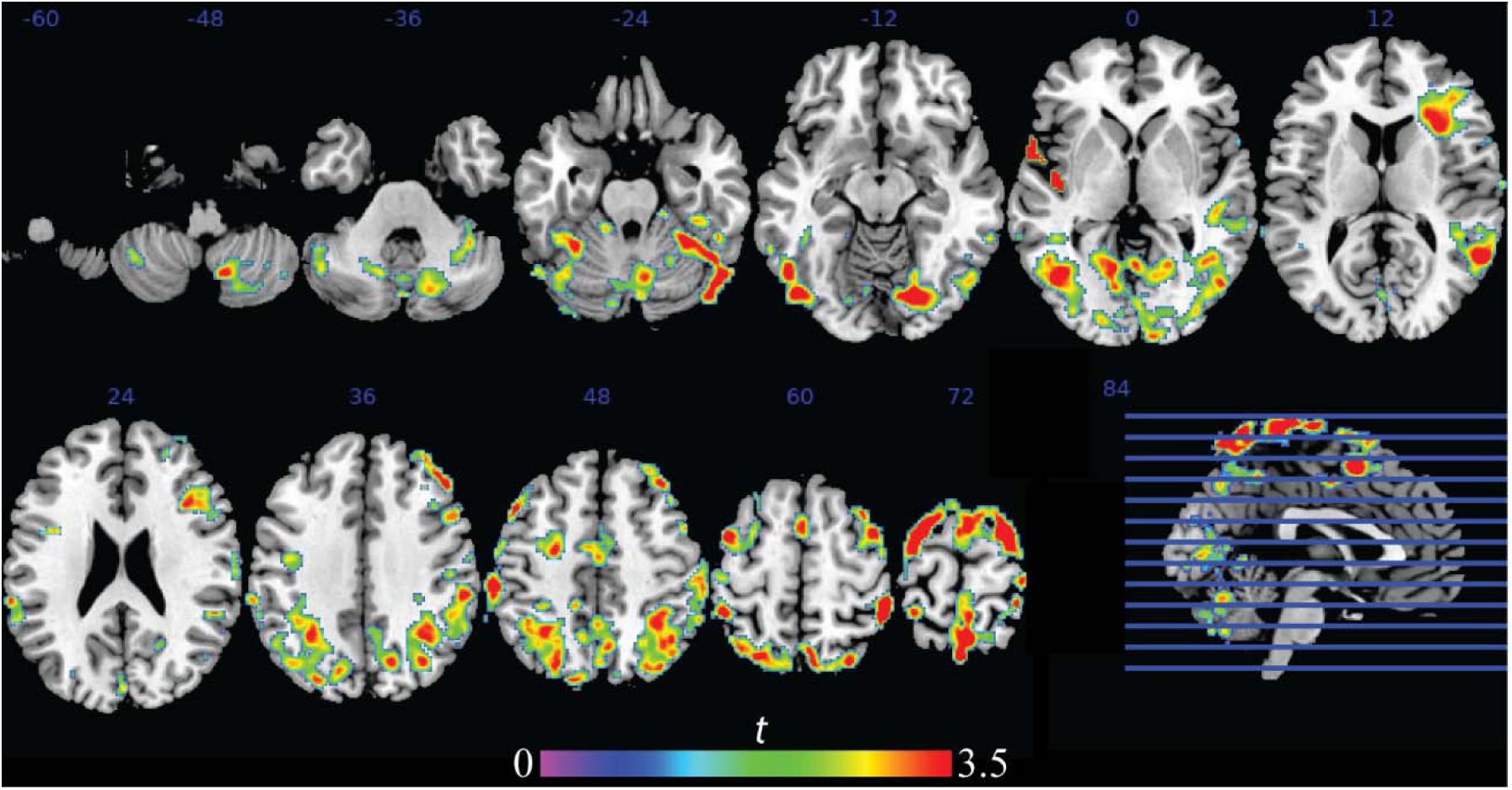
Brain activation patterns to feedback cues during learning within OA.

**Fig. 5.**
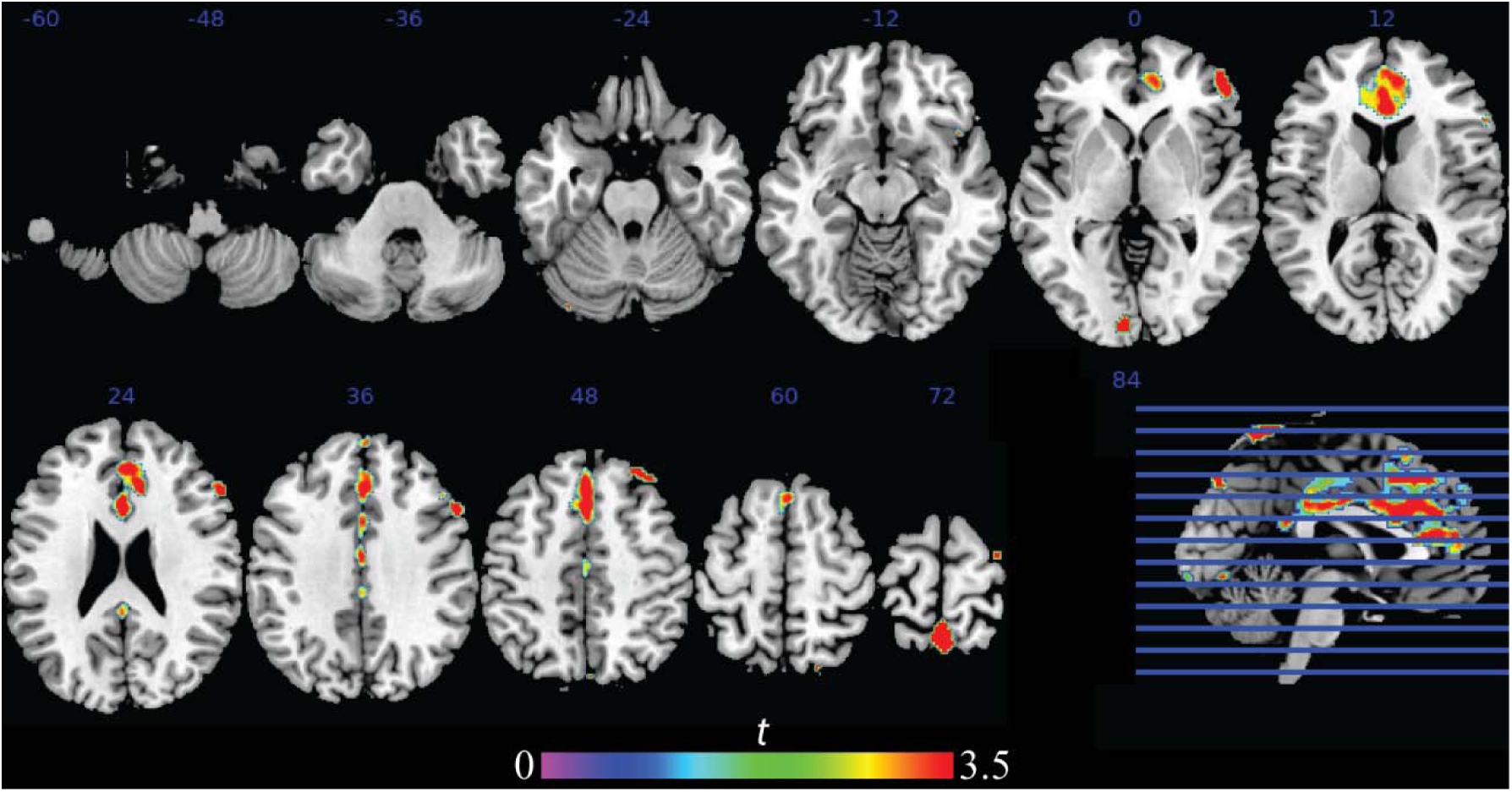
Brain activation patterns to feedback cues during learning within YA.

**Fig. 6.**
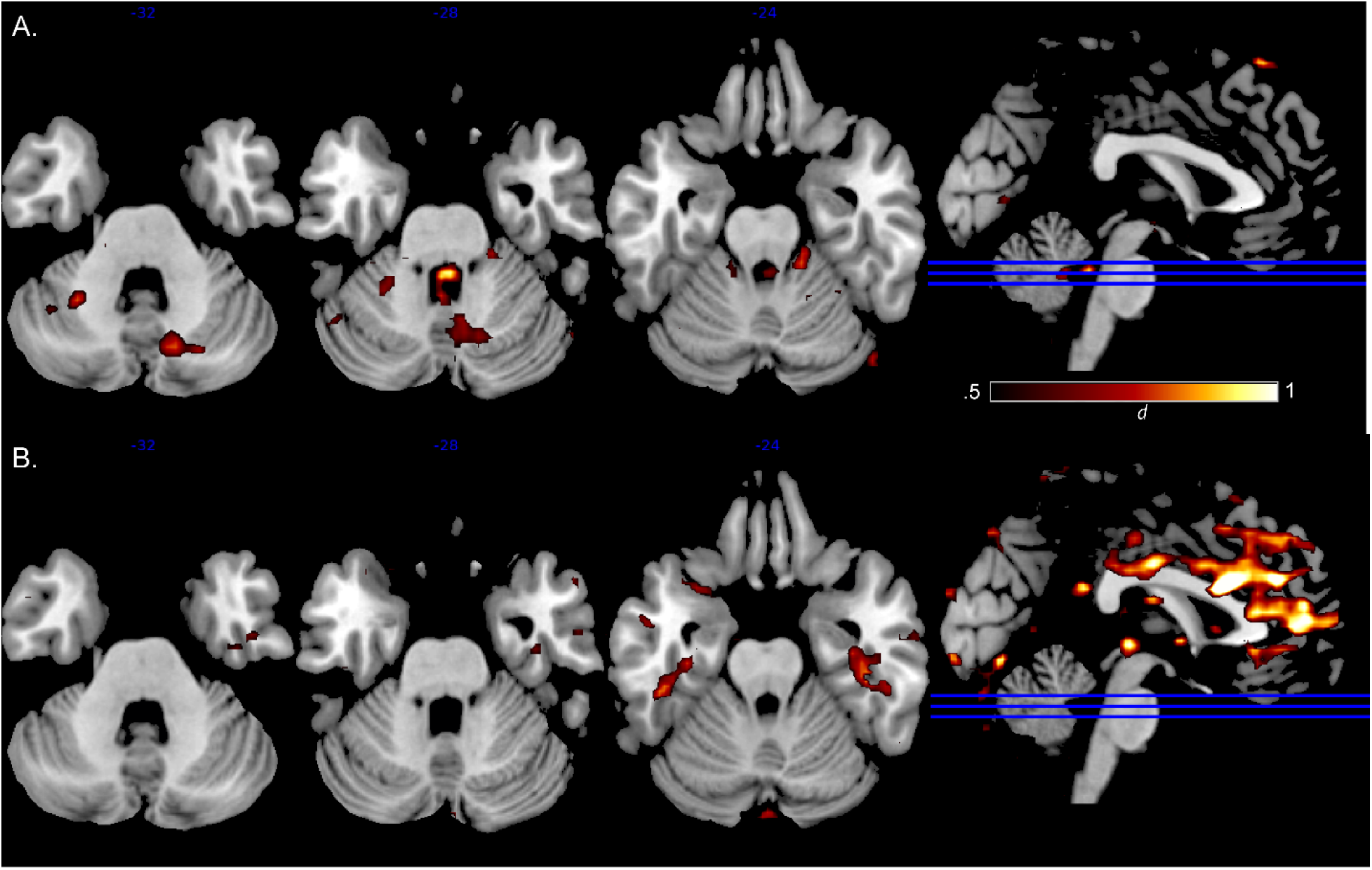
Effect size maps to feedback cues during learning for (A) OA > YA and (B) YA > OA.

**Table 3.**
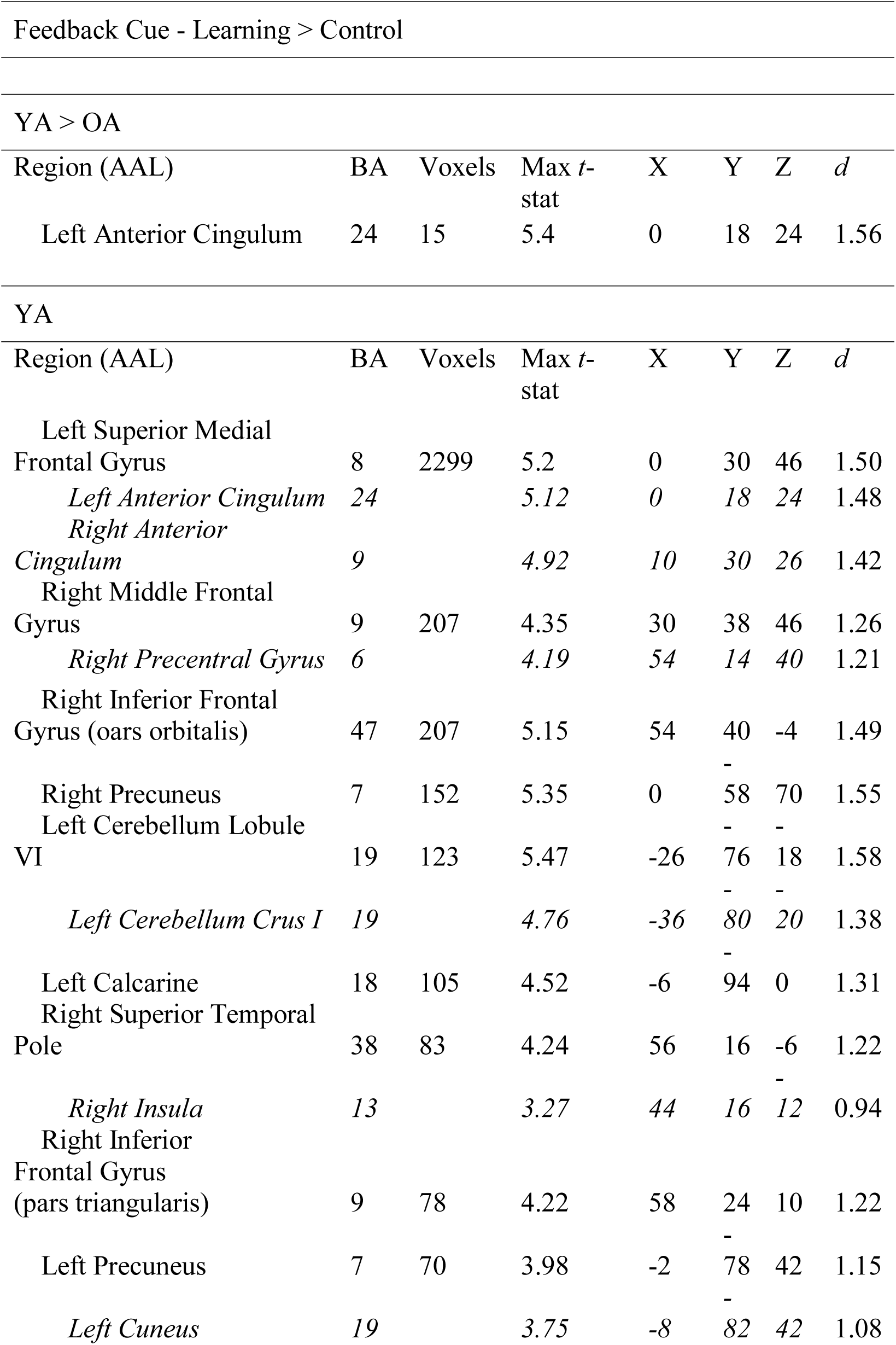
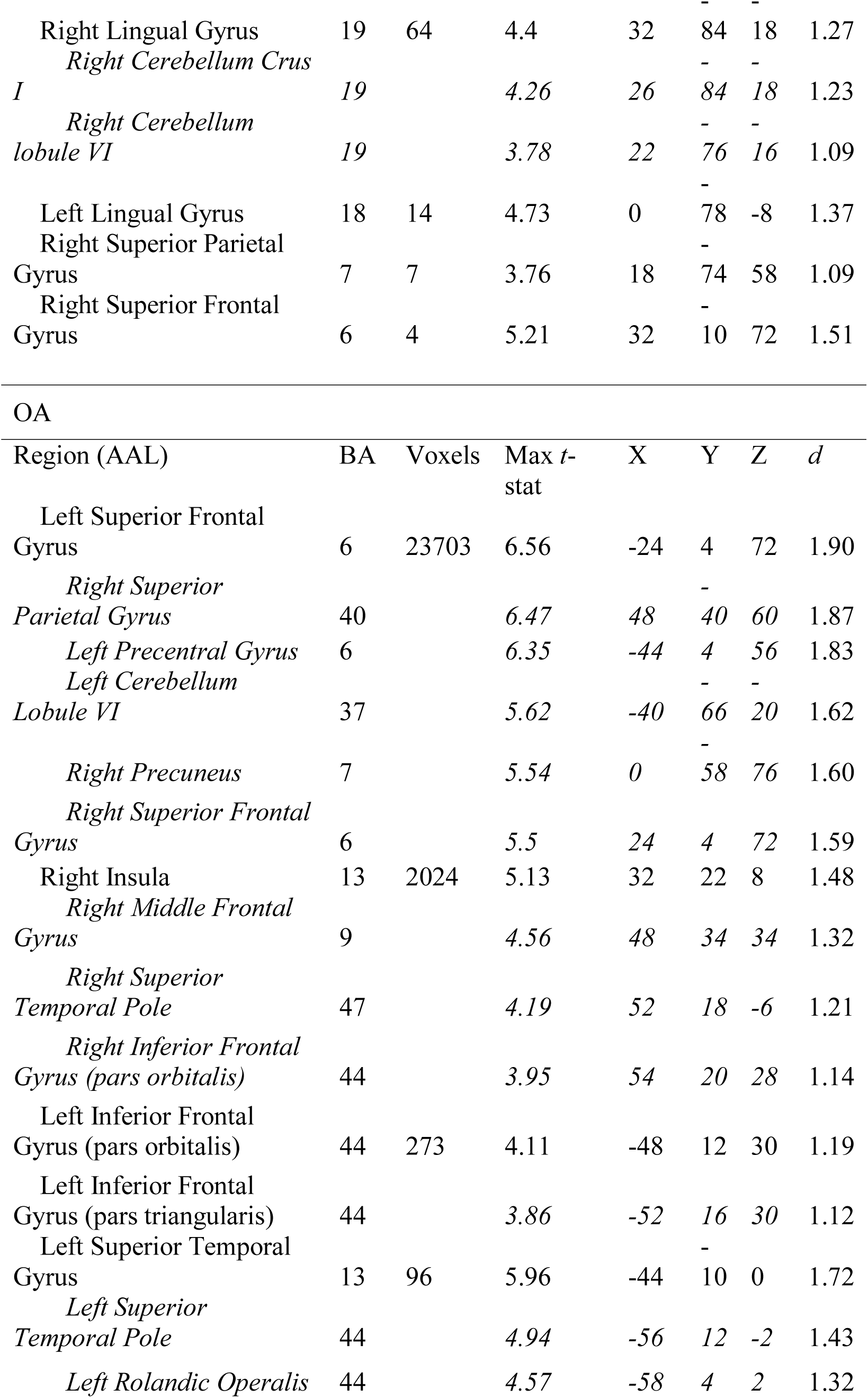
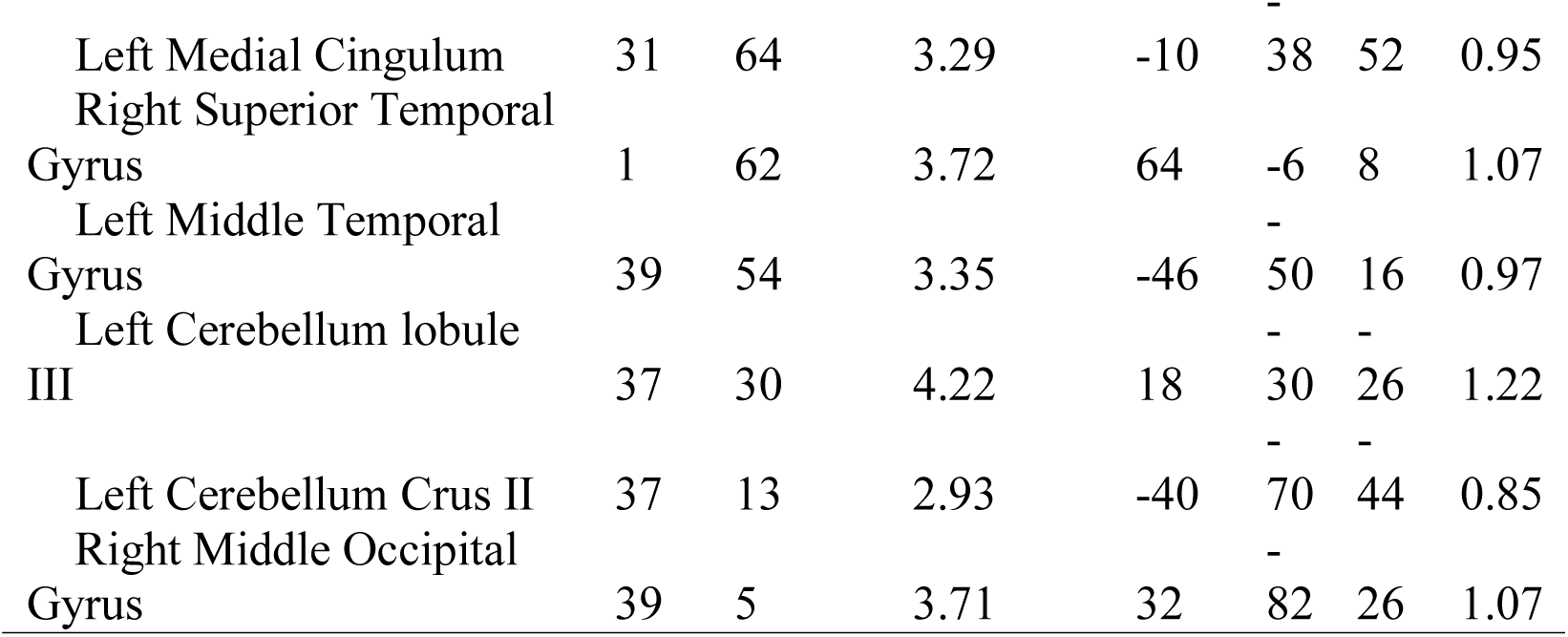
Areas of peak activation in response to feedback cues on correct trials during all learning blocks compared to control blocks. Groups differences and within group data are presented. Regions in italics are local maxima within a larger cluster.

**Table 4.**
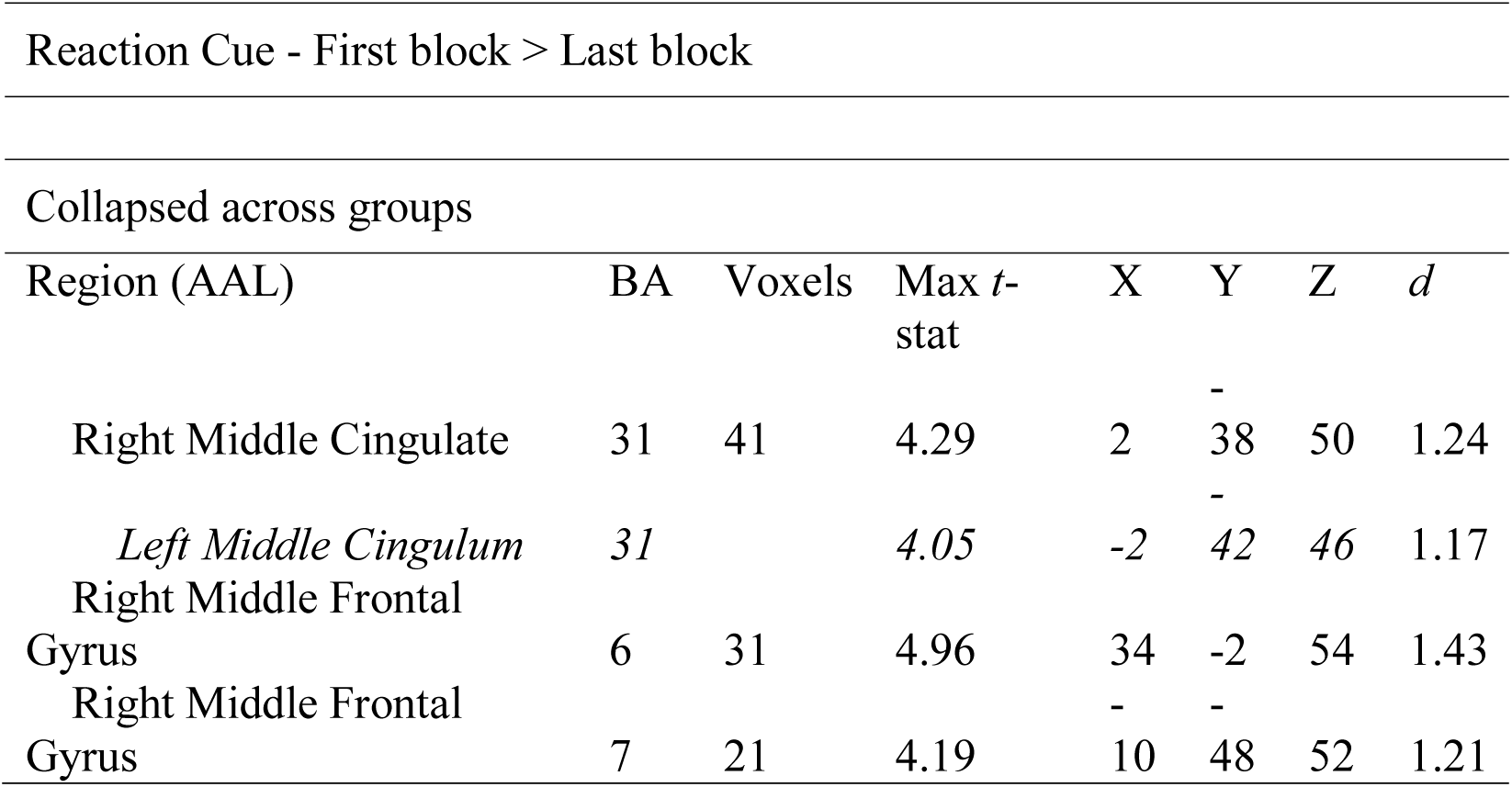
Areas of peak activation in response to reaction cues on correct trials during the first learning block compared to the last learning block. Regions in italics are local maxima within a larger cluster.

#### Block 1 > Block 4

No significant clusters were found in response to instruction cues in any planned or *post-hoc* contrast.

Reaction cues elicited no significant activations in either the group-level or group contrast results. An exploratory analysis revealed bilateral middle cingulum, right frontal middle gyrus, and left precuneus activated when collapsing across groups (Table 3; Figure 7). Effect size calculations indicated OA showed no activation compared to YA in the posterior cerebellum, while YA showed robust posterior cerebellar activation compared to OA, with medium to large effect sizes (Figure 8).

**Fig. 7.**
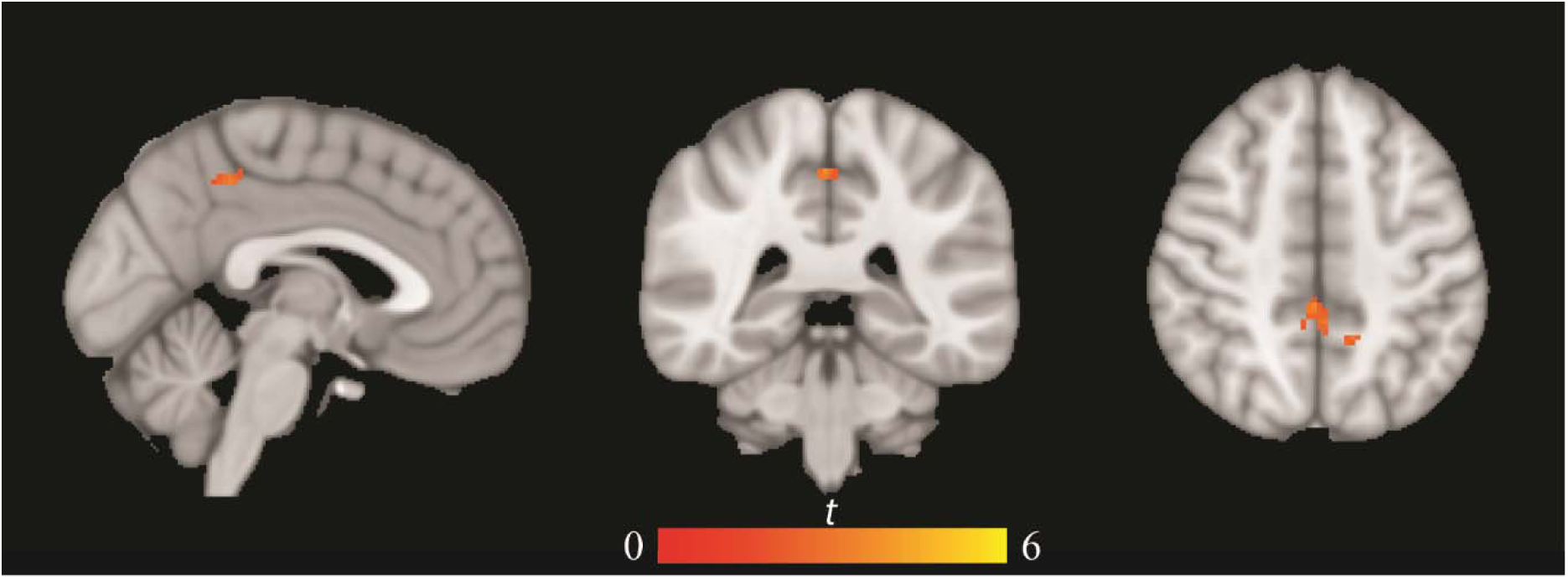
Exploratory analysis showing brain activation patterns to reaction cues during early learning (block 1 > block 4), collapsed across groups.

**Fig. 8.**
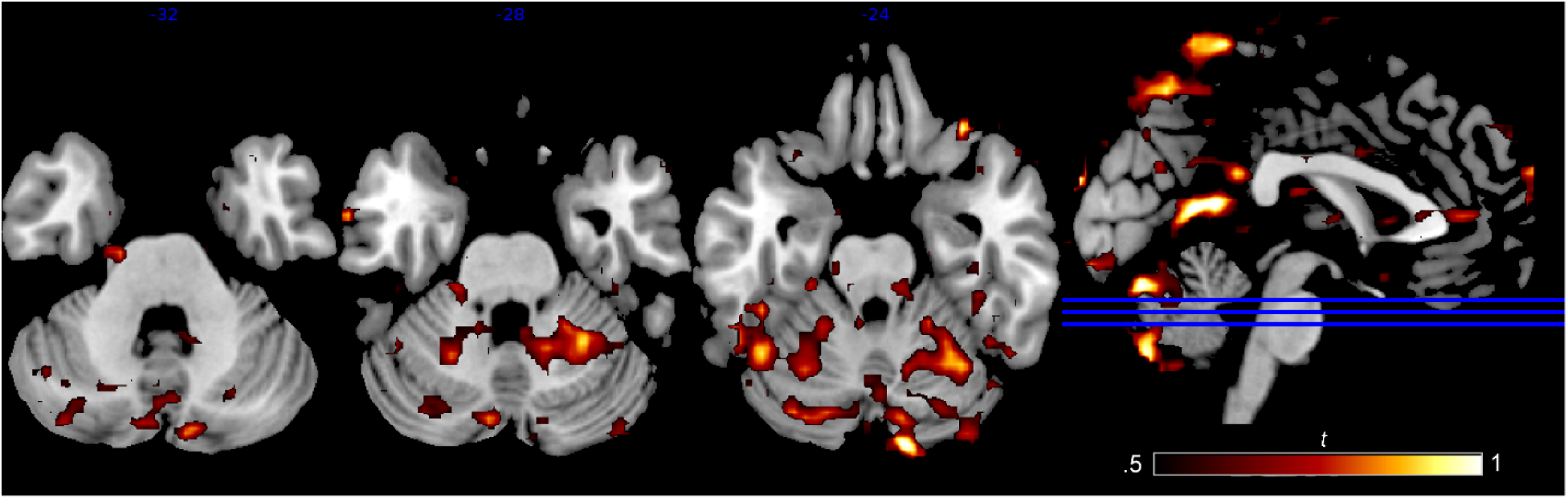
Effect size maps to reaction cues during early learning for (A) OA > YA and (B) YA > OA.

No significant clusters were found in response to feedback cues in any planned or *post-hoc* contrasts.

## Discussion

Here, we adapted an fMRI learning task designed to target both the cerebellum and prefrontal cortical regions, and separate motor planning and responses from second-order cognitive rule learning to investigate CTCC engagement and processing in advanced age (Balsters et al., 2013). Though we are interested in understanding cerebellar functioning in OA, we took a whole brain approach given the importance of CTCC networks to performance. Behavioral performance was largely in line with our hypotheses and previous findings in the older adult higher-order learning and motor learning literatures (Anguera et al., 2011; Bennett, Howard, & Howard, 2007; Bennett, Madden, Vaidya, Howard, & Howard, 2011; Bo et al., 2012; Dennis, Howard, & Howard, 2006; Yassa et al., 2011): OA were outperformed by YA during every learning blocks. Further, while YA were experiencing ceiling effects by block 2, OA only reached an average of 70% correct by the end of block 4. The learning slope analysis expanded on these findings to show that not only did OA perform more poorly than YA in each block, they learned more slowly by comparison.

With respect to the posterior cerebellum, we found that OA showed widespread activation to instruction cues in the learning (greater than control) blocks with a local peak at right lobule VI, a structure thought to be involved in cognitive and emotional processing (Stoodley, 2012; Stoodley et al., 2012). This large cluster also included Crus I and II regions, though local peaks were not present. Within YA, bilateral Crus II, right Crus I, right lobule VI, and right lobule VIII were activated. Of note, Balsters and colleagues (2012) also reported activity in Crus I, Crus II and lobule VI in YA using this paradigm. Taken together, the large cluster in OA is perhaps indicative of increased cerebellar recruitment by OA, though contrast analyses did not show significant group differences. The effect size analysis also supports this interpretation, as OA showed more widespread activation compared to YA (Figure 3). Therefore, while we cannot definitively state that the OA sample differed from YA in cerebellar recruitment while viewing instruction cues, qualitatively, the patterns of recruitment differ.

Cerebellar activity to feedback cues in learning (greater than control) blocks showed left Crus II and lobule VI activation in OA, as well as lobule III, primarily a motor region (Grodd, Hülsmann, Lotze, Wildgruber, & Erb, 2001). While Balsters et al. (2012) did not analyze activity related to feedback cues, activation in Crus I and lobule VI are in line with the findings that these regions are involved in error-related cognitive processes supporting feedforward internal prediction models of cerebellar function (Ben-Yehudah, Guediche, & Fiez, 2007; Ito, 1993, 2008). Additionally, activation of lobule III can perhaps be interpreted as compensation for decreased processing efficiency in the posterior cerebellum, and is in line with arguments that working memory may be supported by the motor system. (Marvel, Morgan, & Kronemer, 2019). YA showed cerebellar activation in response to feedback cues, again with bilateral lobule VI and Crus I activation. In sum, OA showed activation to feedback cues during learning (greater than control) blocks in the posterior cerebellar regions similar to that of YA, in contrast to both the compensatory (Filip et al., 2019; Park & Reuter-Lorenz, 2008) or under activation (Bernard & Seidler, 2014; Bo et al., 2011) hypotheses. Effect size analyses however show OA have more activation in the medial posterior cerebellum than YA, though this is not a significant finding and does not serve to statistically differentiate the age groups (Figure 6).

Outside of the cerebellum, we found that OA, compared to YA, showed increased activity to instruction cues presented in learning (greater than control) blocks in the angular gyrus of the left inferior parietal lobe, a region associated with spatial reasoning, recollection, and recognition (Seghier, 2013). This is consistent with Balsters et al. (2012) and previous findings showing activation in this region to working memory paradigms (Emery et al., 2008). We also found OA showed reduced activity compared to YA to feedback cues in learning (greater than control) blocks in the left anterior cingulate, a task-positive region associated with learning and error correction (Brown, 2005). This may indicate that OA differ in their ability to process feedback and correct for errors, potentially contributing to the slower rates of learning seen in the OA sample. Interestingly, the angular gyrus region is part of the DMN, shown by Andrews-Hanna et al. (2007) to be disrupted in OA. Increased activation in OA compared to YA may suggest reduced suppression of the DMN in OA, consistent with past work predicting increased DMN activity in OA relative to YA (Andrews-Hanna et al., 2007; Damoiseaux et al., 2008; Grady et al., 2010; Grady, Sarraf, Saverino, & Campbell, 2016). Taken together, the results presented here are compatible with previous work suggesting OA show an inability to correctly suppress the DMN during cognitive processing (Chand, Wu, Hajjar, & Qiu, 2017; Grady et al., 2010, 2016; Raichle, 2015) and a seemingly reduced capacity to efficiently bring task-positive, error-correcting and attention-controlling brain regions online to update future responses (Langenecker & Nielson, 2003; Zhu, Zacks, & Slade, 2010). In line with these results, accuracy analyses showed OA are on par with YA by the second control block (a purely first-order rule task), yet do not ever meet YA performance in second-order learning blocks. This perhaps can be explained by the lack of efficient processing of the negative feedback to incorrect responses (as indexed by decreased anterior cingulate activation), resulting in slower learning rates.

Early learning, as indicated by the block 1 greater than block 4 analyses, showed little in terms of activation in either group, though collapsing across groups showed activation to reaction cues in regions associated with working memory and attention. Interestingly, the effect size calculations showed much more widespread cerebellar activation in YA compared to OA, with OA showing no cerebellar recruitment beyond that of YA (Figure 8). While qualitative, this suggests YA were better able to utilize the second-order reaction cue during early learning, whereas OA are less able to incorporate the second-order information.

This study adds to a wealth of knowledge regarding the aging brain. Critically, we demonstrated that contrary to our hypotheses and suggestions from the literature (Bernard & Seidler, 2014; Fillip et al., 2019) there are limited statistically significant differences between OA and YA in terms of cerebellar activation during second-order rule learning, yet qualitatively these groups showed wide differences in recruitment patterns, and effect size analyses suggest that there are substantial group differences of medium to large magnitude. It is important to note the limitations of this study. First, comparing OA to YA is useful in gaining perspective into differences in brain and behavior with age, but analyses fall short of being able to describe the decline across the lifespan. Longitudinal studies would help elucidate those changes and provide insight into whether brain changes predict behavioral changes or vice versa. Second, the learning task was relatively short, and OA never neared 100% accuracy. While we did not see any activation differences in block 1 as compared to block 4, this may be due to the short time frame of the learning for the OA. While we chose to only include 4 learning blocks due to the potential of fatigue in OA, future studies may choose to lengthen the task and/or increase task difficulty for YA in order to more accurately compare learning processes. Additionally, though we have reasonable sample sizes in the context of the aging literature, the power to detect group differences after the appropriate multiple comparison corrections may be to blame for the lack of group contrast effects. This concept is bolstered by the robust differences in effect size images, which show differences that are sub-threshold. Future studies should strive to include larger samples in order to account for this finding.

In this study, we compared the functional activation patterns of a healthy OA sample to a YA sample during a second-order rule learning task to investigate differences in brain and behavior as learning occurs. We found that OA were less able to efficiently learn second-order rules and are qualitatively, but not statistically, different than YA in the ability to bring cognitive regions of the cerebellum online during second-order rule learning. We also suggest that under-activation of the cingulate cortex in OA may be a potential mechanism by which learning is slowed, and replicated previous findings suggesting decreased suppression of the DMN in response to cognitive tasks (Andrews-Hanna et al., 2007; Grady et al., 2010, 2016; Raichle, 2015). This research serves to further detail age-related differences in brain behavior and performance during higher-order cognitive tasks and may help inform future behavioral interventions geared towards improving OA performance.

## Acknowledgements

The authors acknowledge the Texas A&M University Brazos HPC cluster that contributed to the research reported here. <brazos.tamu.edu> J.A.B. was supported in part by a NARSAD Young Investigator Grant from the Brain and Behavior Research Foundation as the Donald and Janet Boardman Family Investigator.

