## Supplemental Figure 1, Supplemental Figure 2 for "Cerebellar and Prefrontal-Cortical Engagement During Higher-Order Rule Learning in Older Adulthood"

Supplemental Figures

Fig. 1) Brain activation patterns to instruction cues during learning within OA.

Fig. 2) Brain activation patterns to instruction cues during learning within YA.

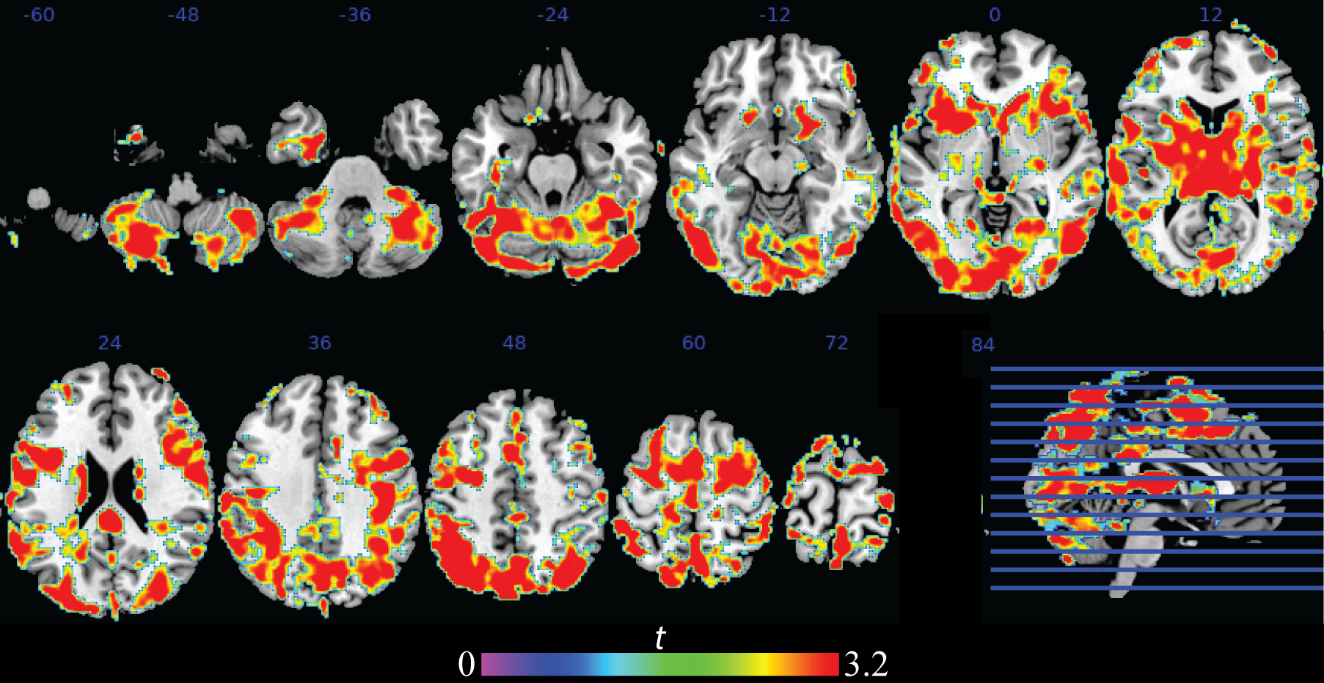

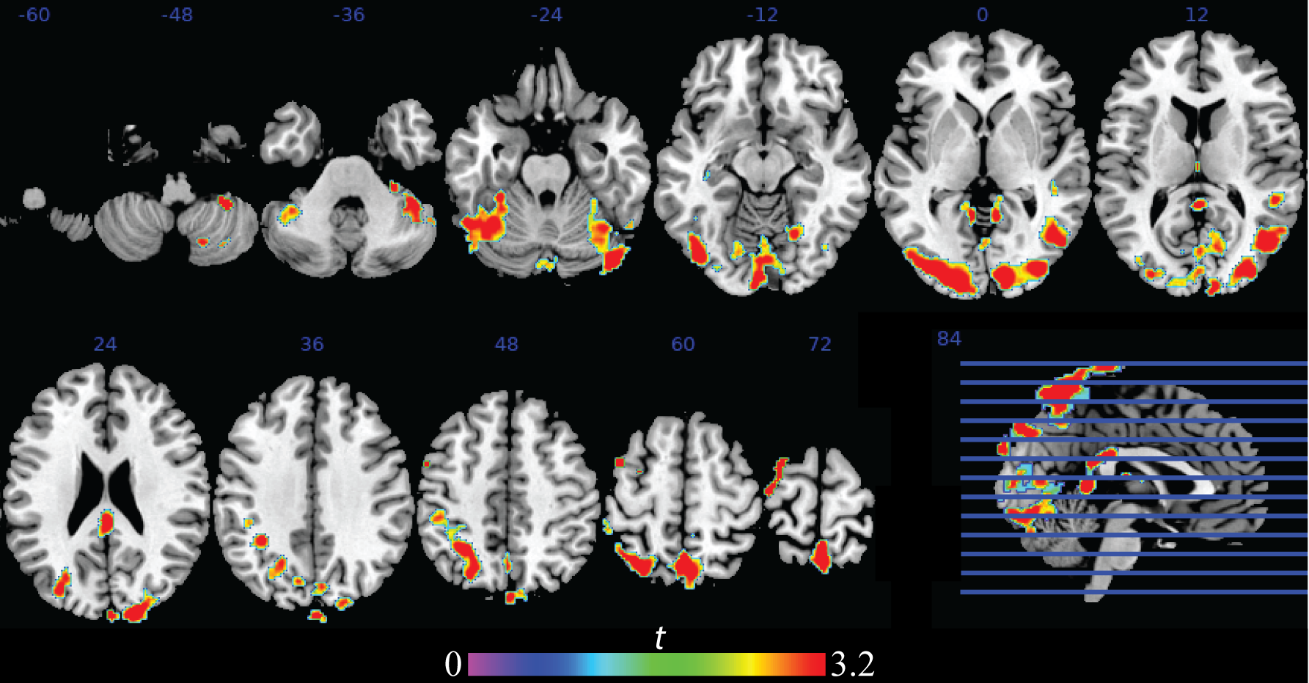
